# Ascending dorsal column sensory neurons respond to spinal cord injury and downregulate genes related to lipid metabolism

**DOI:** 10.1101/2020.05.07.083584

**Authors:** Eric E. Ewan, Oshri Avraham, Dan Carlin, Tassia Mangetti Goncalves, Guoyan Zhao, Valeria Cavalli

## Abstract

Regeneration failure after spinal cord injury (SCI) results in part from the lack of a pro-regenerative response in injured neurons, but the response to SCI has not been examined specifically in injured sensory neurons. Using RNA sequencing of dorsal root ganglion, we determined that thoracic SCI elicits a transcriptional response distinct from sciatic nerve injury (SNI). Both SNI and SCI induced upregulation of ATF3 and Jun, yet this response failed to promote growth in sensory neurons after SCI. RNA sequencing of purified sensory neurons one and three days after injury revealed that unlike SNI, the SCI response is not sustained. Both SCI and SNI elicited the expression of ATF3 target genes, with very little overlap between conditions. Pathway analysis of differentially expressed ATF3 target genes revealed that fatty acid biosynthesis and terpenoid backbone synthesis were downregulated after SCI but not SNI. Pharmacologic inhibition of fatty acid synthase, the enzyme generating palmitic acid, decreased axon growth and regeneration *in vitro*. These results supports the notion that decreased expression of lipid metabolism-related genes after SCI, including fatty acid synthase, may restrict axon regenerative capacity after SCI.

## Introduction

Unlike injured neurons in the peripheral nervous system, which mount a regenerative response and reconnect with their targets, axon regeneration typically fails after spinal cord injury (SCI). In addition to the growth inhibitory environment of the injured spinal cord ^1,2^, gene profiling studies link regenerative failure after SCI to the lack of an intrinsic transcriptional response ^3,4^. This is supported by the conditioning injury paradigm, in which prior injury to the peripheral axons of dorsal root ganglion (DRG) sensory neurons promotes axon regeneration of their central axons after SCI ^5–7^. Peripheral nerve injury upregulates many regeneration-associated transcription factors (RATF) ^8,9^, yet only modest spinal axon regeneration has been observed by overexpressing RATF ^10–12^. This suggests that RATF expression does not sufficiently recapitulate a growth program and that the intrinsic mechanisms underlying regeneration failure after SCI remains incompletely understood.

Whether and how sensory neurons alter their transcriptome after SCI remains unclear because most analyses have used whole DRG after SCI ^3,4,13^, which includes uninjured nociceptors ^14^ and non-neuronal cells ^15^. In this study, we took advantage of a mouse line that labels sensory neurons ascending the spinal cord to unravel their transcriptional response to SCI. We performed RNA sequencing (RNA-seq) experiments from whole DRG and from fluorescence-activated cell sorted (FACS) ascending sensory neurons acutely after SCI and SNI. Our results indicate that ascending sensory neurons alter their transcriptome after SCI, with gene expression changes that differ from SNI. Part of the transcriptional changes observed after SCI are related to a decrease in lipid metabolism and may repress axon regeneration.

## Results

### SCI induces a unique transcriptional response in the DRG compared with PNS injury

Previous studies suggest that transcriptional changes in whole DRG after SCI may occur early after SCI and might not be sustained ^3,4^. SNI was also shown to induce a more robust transcriptional response compared to SCI in whole DRG ^4^. In contrast, SCI induces more changes in 5-hydroxymethylcytosine than SNI, an epigenomic mark that has transcriptional regulatory roles ^13^. These studies suggest that SCI may elicit a transcriptional response in DRG, but a detailed analysis of how SCI affect gene expression in DRG and sensory neurons has not been performed in detail. To better understand the transcriptional response in DRG after SCI and SNI, we performed RNA-seq of L4 DRG one day (1d) after SCI and analyzed our results with a previously generated RNA-seq data set collected 1d after SNI (**Fig 1A**; ^16^). We found that SCI elicits a less robust transcriptional response in the DRG compared to SNI, with fewer differentially expressed (DE) genes compared to SNI (**Fig 1B, Supplementary Table 1,2**). There were few DE genes overlapping between conditions (~15%) and many (~9%) were inversely expressed (**Fig 1B**). KEGG (Kyoto Encyclopedia of Genes and Genomes biological) pathway analysis of DE genes identified different pathways enriched after SNI and SCI (**Fig S1A-B**), suggesting that the SCI response is distinct from the SNI response.

**Figure 1.**
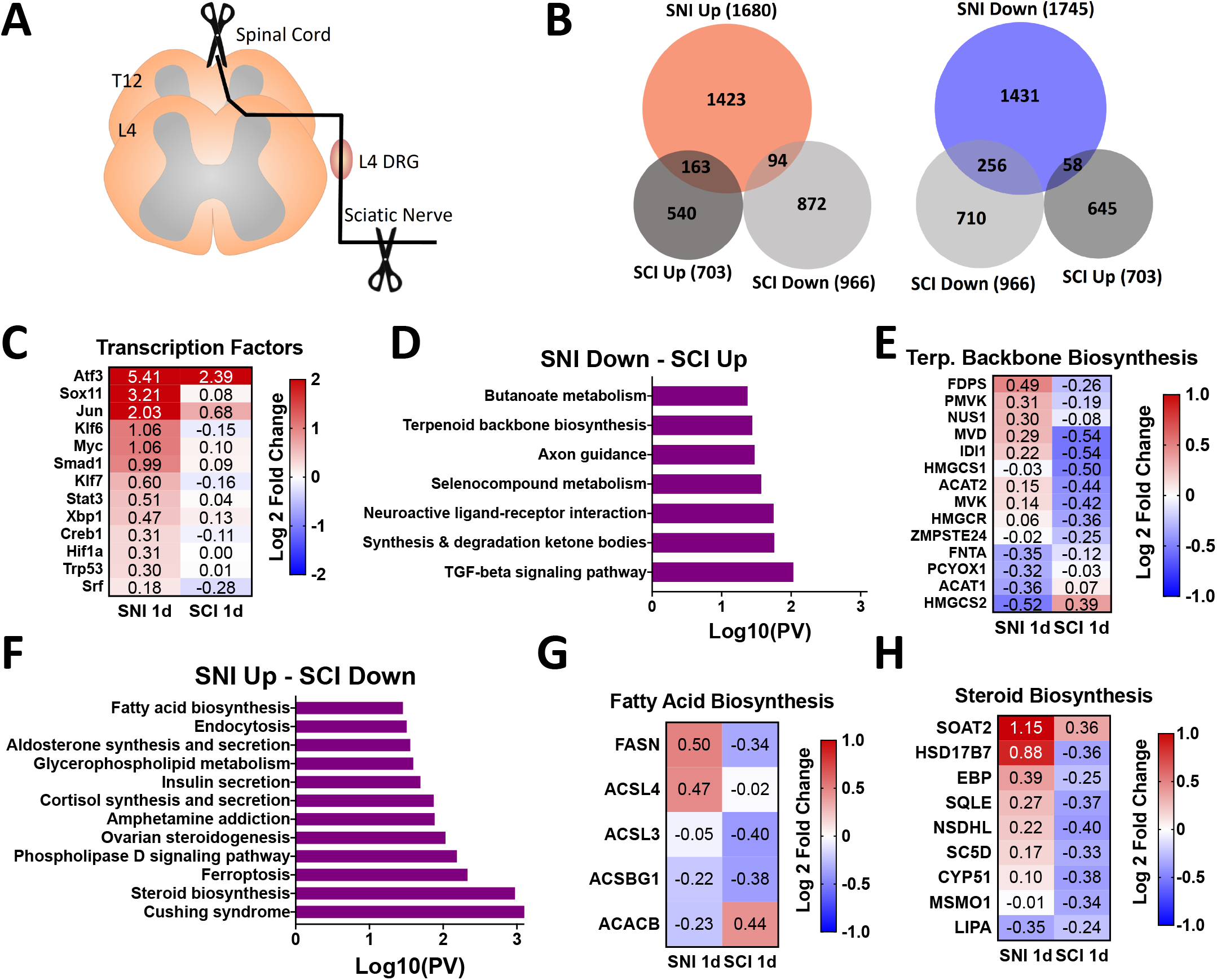
Spinal Cord Injury (SCI) induces an acute transcriptional response in the dorsal root ganglion (DRG) that differs from sciatic nerve injury (SNI). **A**) Schematic of the experimental design for L4 DRG RNA sequencing after SNI and SCI. **B)** Proportional Venn diagrams for differentially expressed (DE) genes upregulated (red) or downregulated (blue) after SNI (n=3 mice) and SCI (n=4 mice) (p-adj < 0.1). **C**) Heatmap of known regeneration-associated transcription factors (RATF’s) after SNI and SCI. **D**) Pathway analysis of DE genes downregulated after SNI and upregulated after SCI (KEGG 2016). **E**) Heatmap of genes associated with terpenoid backbone biosynthesis after SNI and SCI. **F**) Pathway analysis of DE genes upregulated after SNI and downregulated after SCI (KEGG 2016**). G-H**) Heatmap of genes associated with fatty acid biosynthesis and steroid biosynthesis after SNI and SCI.

We next examined the expression of RATFs, given their important roles in establishing a regenerative axon growth program ^9,17^. Whereas all RATFs examined were upregulated after SNI, only ATF3 and Jun were increased after SCI (**Fig 1C, Supplementary Table 3**). The activation of RATFs is believed to arise in part from retrograde transport of kinases from the injury site back to the cell soma ^18,19^. The dual leucine zipper kinase DLK is required for retrograde injury signaling and induction of RATF after nerve injury ^16,20^, and is significantly decreased after SCI (0.29 fold, p-adj < 0.05). To determine if the limited transcriptional upregulation of RATF’s after SCI relates to decreased levels of DLK, we used previously generated RNA-seq data that examined the transcriptional response to nerve injury in mice lacking DLK (DLK KO; ^16^). Comparison of DE genes and KEGG pathway analysis in DLK KO following SNI and wildtype mice following SCI revealed little overlap between conditions (**Fig S1B-D**), suggesting that the diminished RATF upregulation after SCI is unlikely mediated by reduced DLK signaling.

The unique transcriptional response of SCI in the DRG compared to SNI may result in gene expression that actively represses axon regeneration after SCI. To examine this possibility further, we performed KEGG pathway analysis on DE genes that were inversely expressed between conditions. We found that inversely expressed DE genes were associated with biosynthetic and lipid metabolic pathways (**Fig 1D-H, Supplementary Table 4**). These pathways were also some of the most enriched by SCI when all downregulated DE genes were examined (**Fig S1B**). This is particularly interesting since previous studies using DRG cultures revealed that synthesis of phosphatidylcholine and cholesterol is required for axonal growth ^21–23^, although more recent studies suggest that decreasing cholesterol levels promote axon growth ^24,25^. Furthermore, in the optic nerve injury model it was shown that increasing phospholipid synthesis and decreasing triglyceride synthesis promotes axon regeneration ^26^. Together, these results suggest that defects in lipid synthesis may restrict axon regeneration after SCI.

### ATF3 and Jun upregulation after SCI is not sufficient to promote growth *in vitro*

ATF3 and Jun are required for peripheral nerve regeneration ^27 28^. When co-expressed in cultured DRG neurons, ATF3 and Jun promote axon growth *in vitro* ^29^ and regeneration of the central axon branch of sensory neurons *in vivo* ^30^. We thus determined if ATF3 and Jun upregulation after SCI occurs in ascending sensory neurons. We used Thy1-YFP16 mice ^31^, which label large diameter ascending sensory neurons ^32,33^. Unlike small diameter nociceptors, these sensory neurons ascend the spinal cord and are directly injured by thoracic SCI ^34^ (**Fig 2A**). Indeed, all low-threshold mechanoreceptors ascend the entire dorsal column up to cervical level, and nearly all proprioceptors ascend 6-8 segments along the spinal cord ^34^. Ascending sensory neurons are also distinct from nociceptors in that they express NF200 (*Nefh*) but not TrkA ^35^ and we confirmed that YFP neurons were NF200 positive and TrkA negative (>90%, **Fig 2B-C**). We found that after SNI, ATF3 and Jun were expressed in nearly two-thirds of all DRG neurons, with equal numbers between YFP positive and negative neurons (**Fig 2D-E; Fig S2A-B**). In contrast, only ~5% of all DRG neurons expressed ATF3 and Jun after SCI, but ~20% of YFP neurons were ATF3 and Jun positive after SCI (**Fig 2D-E; Fig S2A-B**). We next asked whether this increased ATF3 and Jun expression in YFP neurons promotes axon growth *in vitro* 3 days after SCI. We observed that unlike SNI, SCI failed to induce a conditioning effect *in vitro* in YFP neurons compared to naive (**Fig 2F-G**). These results suggest that the other transcriptional changes elicited by SCI we observed (**Fig 1D-H**) may repress the pro-growth effects associated with ATF3 and Jun expression. It is also possible that ATF3 and Jun need to be expressed in the same cells to synergize ^29^. We think this is likely, but limitations with antibodies did not permit us to confirm co-expression of ATF3 and Jun in injured DC neurons.

**Figure 2.**
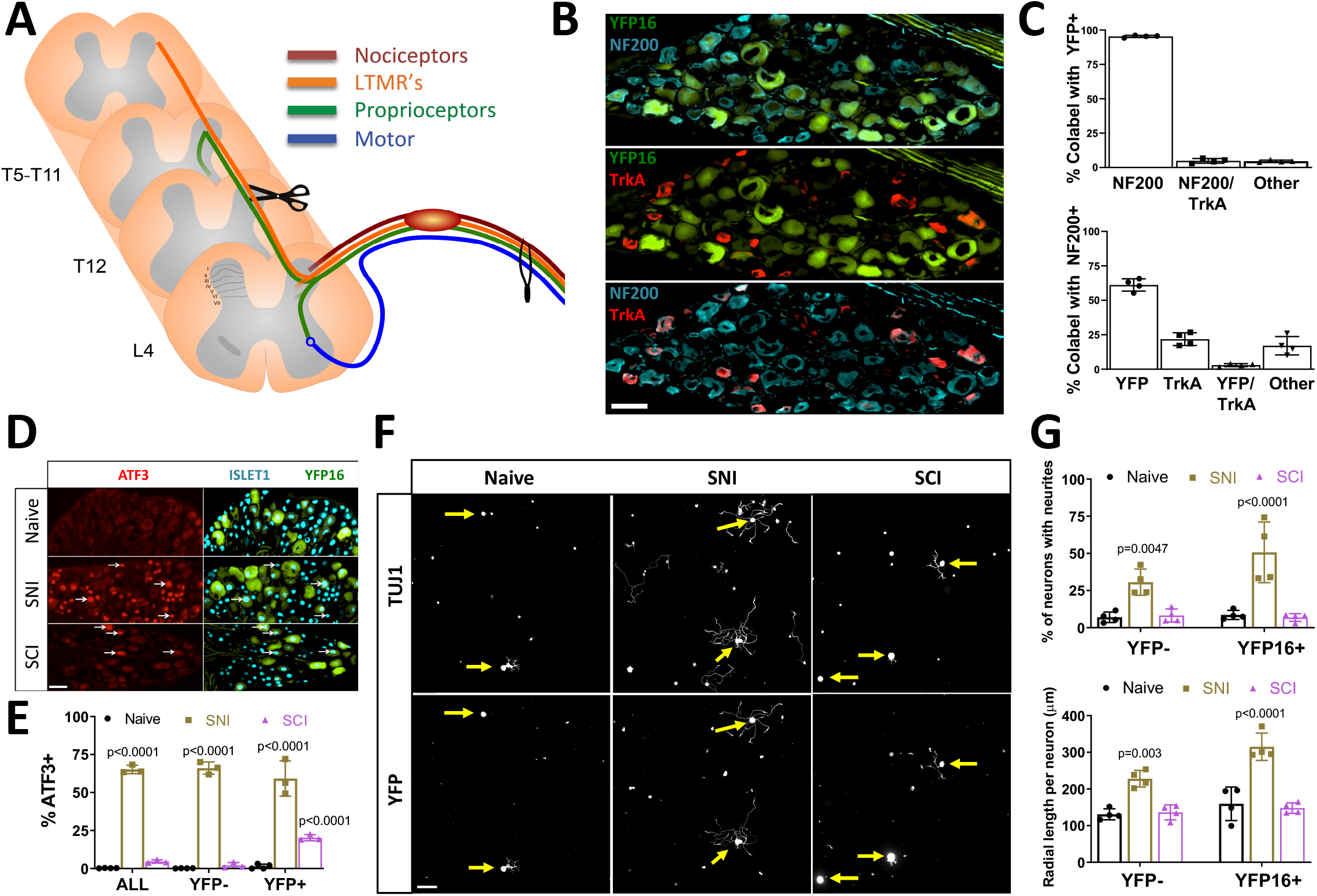
Spinal Cord Injury (SCI) does not induce a pro-growth state in ascending sensory neurons *in vitro*. **A**) Schematic of the experimental design and peripheral and central projects of major sensory neuron subtypes of L4 dorsal root ganglion (DRG) neurons. **B**) Representative images of L4 dorsal root ganglion (DRG) neurons from Thy1YFP16 mice labeled with NF200 (blue) and TrkA (red) antibodies. **C**) Quantification of **B** indicating percent overlap of YFP, NF200, and TrkA labeling in Thy1YFP16 mice (n=4 mice). **D**) Representative images of L4 DRG neurons labeled with ATF3 and Islet1 antibodies in Thy1YFP16 mice in naive or 3 days after sciatic nerve injury (SNI) or SCI. **E**) Quantification of **D** indicating percentage of ATF3 positive, Islet-1 labeled neuronal nuclei in all neurons, as well as YFP negative and YFP positive neurons, for each condition (n=3 mice/group; 2-way ANOVA). White arrowheads point to ATF3 positive neuronal nuclei. **F**) Representative images of cultured cells from L4 DRG labeled for all neurons (TUJ1) and YFP positive neurons in naïve or 3 days after SNI or SCI. **G**) Quantification of **F** indicating the percentage of neurons extending neurites and the average radial length of neurites, in YFP negative and YFP positive neurons, for each condition (n=4 mice/group; 2-way ANOVA). Yellow arrows point to the cell body of YFP positive neurons.

### SCI induces a unique transcriptional response in DC neurons compared to SNI

Neurons are outnumbered by non-neuronal cells at least 10-fold in the DRG ^15^ and not all lumbar DRG neurons ascend the spinal cord and are directly injured by T9 SCI ^34^. Thus, to define the transcriptional changes elicited by SCI in ascending sensory neurons specifically, we performed RNAseq 1 and 3 days after SCI or SNI in FACS-sorted ascending sensory neurons from YFP16 mice (**Fig 3A**). This approach allowed us to enrich for ascending sensory neurons, based on the expression levels of known neuronal marker genes ^35^ when comparing FACS-sorted and whole DRG samples (**Fig 3B, Supplementary Table 3,5**). We found that SCI elicited fewer DE genes compared to SNI at 1d and 3d (**Fig 3C-D, Supplementary Table 6,7**). There was little overlap between the two conditions, with only ~37% for upregulated genes and ~23% for downregulated genes (**Fig 3C**). Whereas the numbers of DE genes were increased from 1 to 3 days after SNI, DE genes decreased from 1 to 3 days after SCI (**Fig 3D, Supplementary Table 6,7**), suggesting that the SCI response is not sustained over time. This may explain why few transcriptional changes in whole DRG were observed 7 days after SCI ^3^.

**Figure 3.**
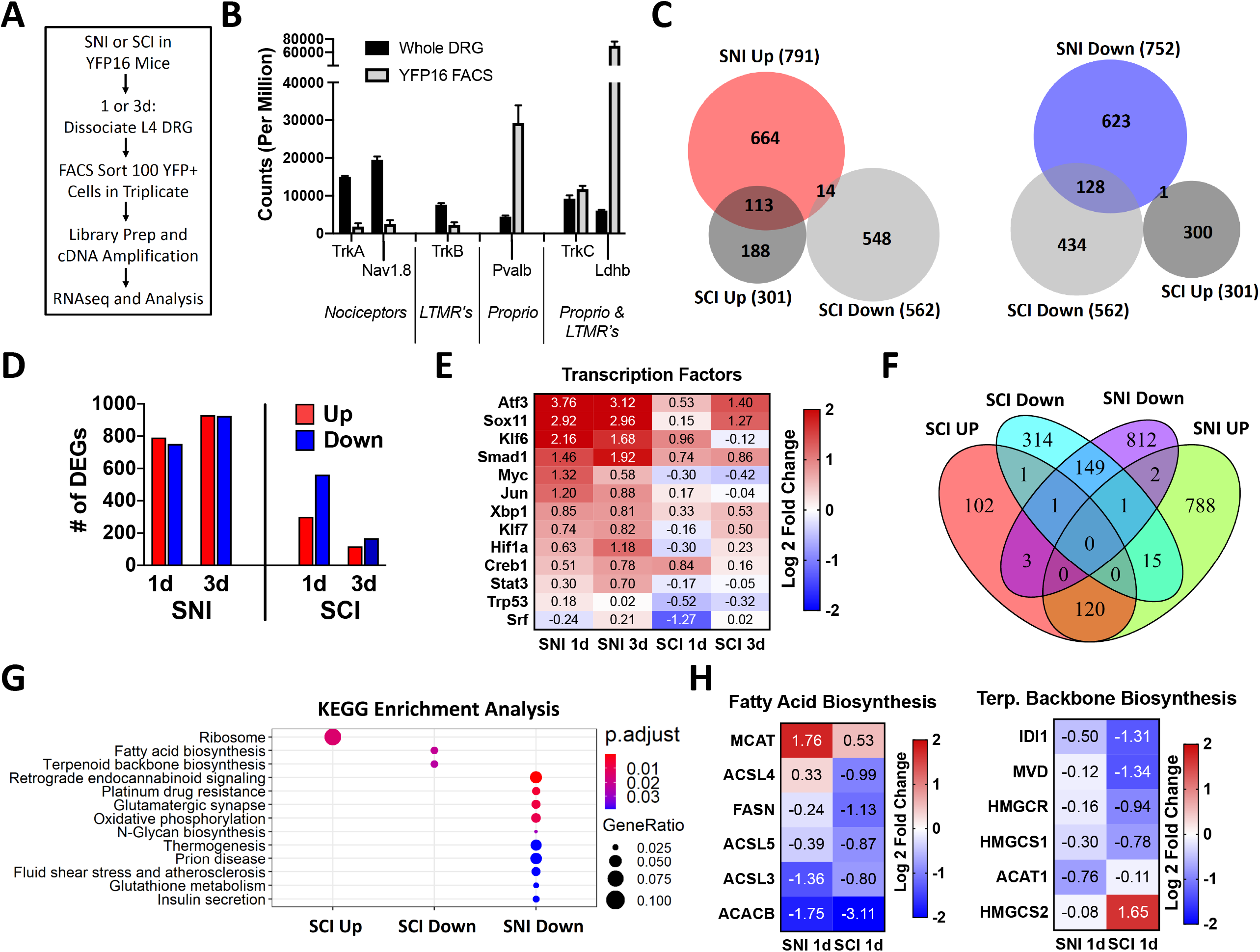
Spinal Cord Injury (SCI) induces an acute transcriptional response in ascending sensory neurons that mostly differs from sciatic nerve injury (SNI). **A**) Schematic of the experimental design for fluorescence-activated cell sorting (FACS) of ascending sensory neurons from Thy1YFP16 mice 1 or 3 days after SNI or SCI (n=3 mice per group, except n=4 mice for naïve and SCI 1d). **B**) Total counts of genes associated with three predominant neuronal subtypes in the DRG (nociceptors, low-threshold mechanoreceptors (LTMR), and proprioceptors (Proprio) (**C**) Proportional venn diagrams for differentially expressed (DE) genes upregulated (red) or downregulated (blue) 1 day after SNI and SCI in ascending sensory neurons, and **D**) Number of DE genes 1 and 3 days after SNI or SCI (p-adj < 0.1, r = 2 for RUVr). **E**) Heatmap of known regeneration-associated transcription factors (RATF’s) 1 and 3 days after SNI and SCI in ascending sensory neurons. **F**) Venn diagram and **G**) KEGG enrichment analysis of DE genes 1 and 3 days after SNI or SCI that have ATF3 binding sites. SNI upregulated DE genes did not lead to significant pathway enrichment. **H**) Heatmap of genes that ATF3 binding sites associated with fatty acid biosynthesis and terpenoid backbone biosynthesis 1 day after SNI and SCI in ascending sensory neurons. Only HMGCS2 does not have an ATF3 binding motif.

We next examined the expression of RATF known to promote axon growth ^9,17^. We found that all assessed RATFs were upregulated in ascending sensory neurons 1 and 3 days after SNI (**Fig 3E, Supplementary Table 5**). In contrast, only ATF3 and Smad1 were increased 1 and 3 days after SCI. KLF6 and Creb1 expression increased at 1d, while Sox11 increased at 3d after SCI (**Fig 3E, Supplementary Table 5**). The SNI-induced upregulation of RATFs in ascending sensory neurons was remarkably similar to the response we previously reported in FACS-sorted nociceptors (**Fig S3A-B**) ^36^. This is consistent with recent findings indicating that after peripheral nerve injury most sensory neurons adopt a similar transcriptomic state ^37 28^. The unique effects of SCI on ascending sensory neurons are thus unlikely related to any intrinsic transcriptional differences between sensory neuron subtypes within the DRG. To better understand the transcriptional differences in ascending sensory neurons after SNI and SCI, we next performed KEGG pathway analysis of all DE genes. We identified different pathways enriched after SCI and SNI (**Fig S3C-F**), with the 5 most enriched KEGG pathways associated with biosynthetic and lipid metabolic pathways 1d after SCI (**Fig S3D**). The top 3 downregulated biosynthesis pathways 1d after SCI were steroid biosynthesis, fatty acid biosynthesis, and terpenoid backbone biosynthesis (**Fig S3D**), all of which were also identified in whole DRG analysis 1d after SCI (**Fig S1B**). None of these pathways were downregulated by SNI, and instead steroid biosynthesis was upregulated both 1d and 3d after SNI (**Fig S3C,E**). Interestingly, we found that only 43% and ~13% of DE genes identified in FACS-sorted ascending sensory neurons after SNI and SCI, respectively, overlapped with DE genes identified from whole DRG (**Fig S3G, S3H**). Many genes are likely not identified in whole DRG because unlike SNI, SCI only directly injures the ascending sensory neurons subpopulation within the DRG ^34^.

We next examined ATF3 target genes, because ATF3 is required to induce the transcriptional re-programming of all neuron subtypes after nerve injury, and ATF3 DNA-binding motifs are enriched in the common injury gene set across neuronal subtypes ^28^. We identified DE genes that have ATF3 binding sites in the promoter region using Patser software ^38^ and ATF3 binding matrix obtained from JASPAR database. Because a majority of the shared DE genes identified at 1 and 3 days post injury changed in the same direction under the same injury condition, we combined the genes for downstream analysis. We found a total of 706 unique ATF3 target genes regulated by SCI and 1891 regulated by SNI, with only 289 common genes (**Fig 3F, Supplementary Table 8**). KEGG analysis revealed that fatty acid biosynthesis and terpenoid backbone synthesis were specifically downregulated after SCI but not SNI (**Fig 3G**). Further analysis of the ATF3 target genes in these pathways indicated that most genes were more significantly downregulated after SCI than SNI, and some were upregulated by SNI (**Fig 3H, Supplementary Table 8**). GO analysis for molecular function of ATF3 target genes further revealed little overlap between SCI and SNI, and also revealed that the downregulation of genes related to ion channel, which is believed to represent in part the loss of neuronal identity after injury^28^, only occurs after SNI (**Fig S4**). These results reveal that lipid biosynthetic pathways are downregulated in ascending sensory neurons after SCI and may repress axon regeneration.

### Fatty acid synthase (FASN) inhibition decreases DRG axon growth *in vitro*

FASN is an enzyme responsible for *de novo* fatty acid synthesis ^39^. *Fasn* is one of the genes in the fatty acid biosynthesis pathway containing an ATF3 binding motif and is downregulated in ascending sensory neurons specifically after SCI (**Fig 3H, Supplemental Table 8**). FASN synthesizes palmitic acid, which is the substrate for the synthesis of more complex lipids including phospholipids ^39^. Given the role of lipid synthesis in axon growth ^21,23,40^, we tested whether inhibiting FASN with platensimycin, an inhibitor that was shown to effectively inhibit fatty acid synthesis *in vitro* ^41^, impairs axon growth. Adult DRG neuronal cultures were treated with platensimycin or vehicle control 30 minutes after plating and axon growth was assessed 24 h later. Platensimycin decreased axon growth by ~40% compared to vehicle controls (**Fig 4A-B**). Since adult DRG cultures include both neurons and non-neuronal cells, we cannot rule out the potential impact of FASN inhibition on non-neuronal cells in decreasing axon growth *in vitro.* Indeed, neurite growth in culture hippocampal neurons was shown to be supported by phospholipid-loaded lipoproteins secreted from glial cells ^42^. We also recently showed that FASN is required in satellite glial cells, which completely surround sensory neurons, to promote axon regeneration in peripheral nerves ^43^. Therefore, to test if FASN is required intrinsically in neurons to promote axon growth, we used a spot culture assay, in which embryonic DRG are dissociated and cultured in a spot, allowing axons to extend radially from the spot ^44,45^. By including the mitotic inhibitor 5-deoxyfluoruridine (FDU), we can obtain a pure neuronal culture that does not contain other non-neuronal cell types ^43^. This assay recapitulates for the most part what can be observed *in vivo* in the nerve ^44–46^, and is thus suitable to test compounds affecting axon regeneration. Platensimycin was added to the media 6 days after plating. Axons were cut using a microtome blade on day 7 and allowed to regenerate for 24h, fixed and stained for SCG10 to visualize axon regeneration ^47^. Regenerative axon length was measured from the visible blade mark to the end of the regenerating axon tips. We observed that platensimycin decreased axon regenerative capacity (**Fig 4C-D**). FASN generates palmitic acid, which is used in part to make phospholipid ligands for the peroxisome proliferator-activated receptor (PPAR) family of transcription factors. PPARs are ligand-activated nuclear receptors that bind lipid signaling molecules to transduce the signals derived from the metabolic environment to control gene expression ^48^. In adipose tissue, FASN is required for generating endogenous ligands for PPARγ ^49^ and in neurons, PPARγ contributes to the pro-regenerative response after axon injury ^50^. We found that the PPARγ agonist rosiglitazone partially rescued the axon regeneration defects induced by FASN inhibition (**Fig 4CD**). These results indicate that in neurons, FASN promotes axon growth in part by generating the ligands for PPARγ.

**Figure 4.**
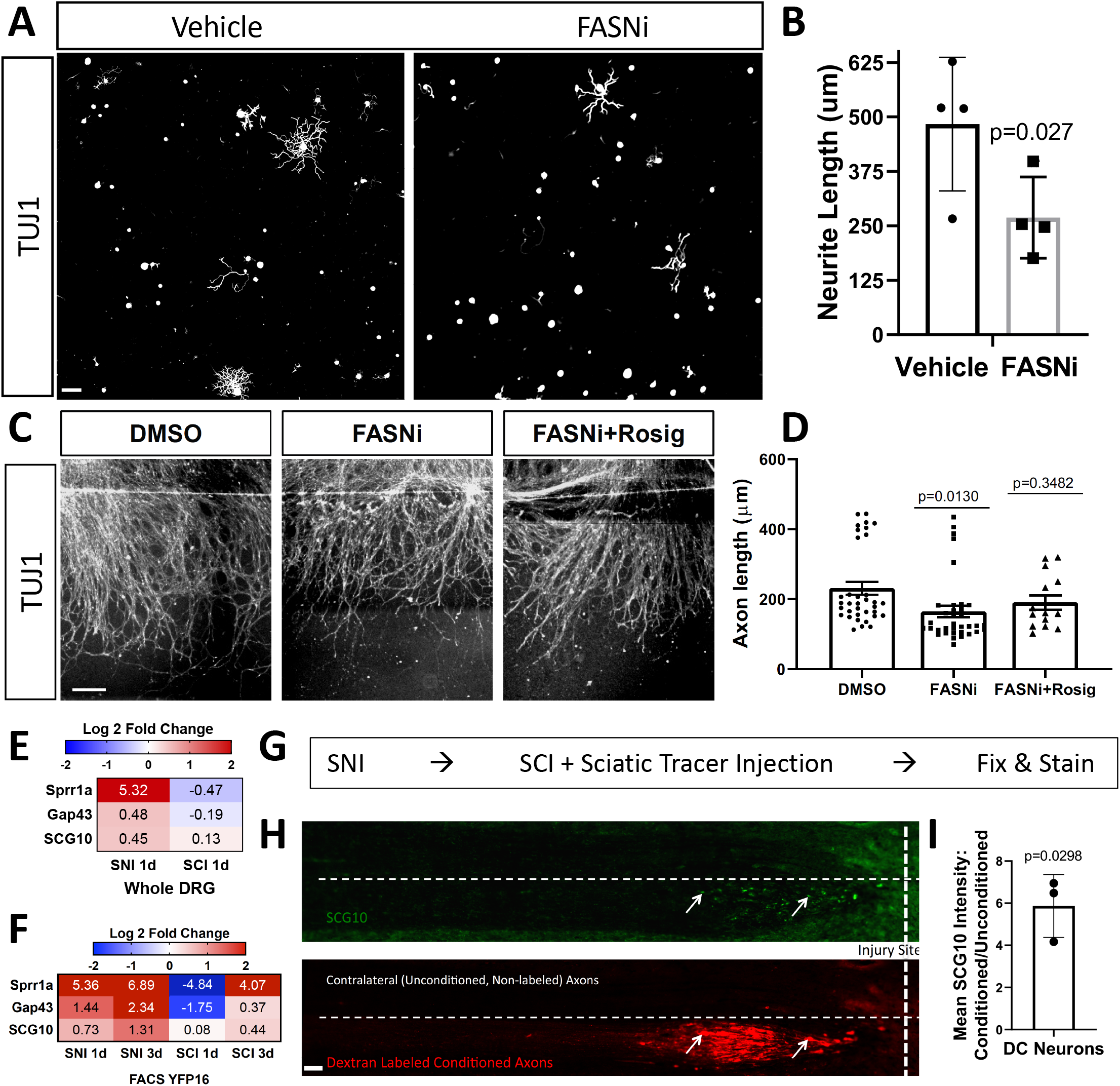
Fasn inhibition decreases sensory neuron growth *in vitro.* **A**) Representative images of TUJ1-labeled sensory neurons cultured from L4 dorsal root ganglion (DRG), that were treated with vehicle or 1um of platensimycin 1 hour after plating, and then allowed to grow for 24hr in culture. **B**) Quantification of **A** indicating the average neurite length for each neuron for each condition; automated neurite tracing and length quantifications were performed using Nikon Elements software (n=4 mice/condition; Unpaired t-test). **C**) Embryonic DRG spot cultures axotomized at DIV7 after a 24 h pre-treatment with DMSO as a control, platensimycin or platensimycin combined with rosiglitazone. Cultures were fixed after 24h and stained with SCG10. Scale Bar: 50 μm. **D**) Quantification of **C** indicating the distance of regenerating axons measured from the injury site. **E,F**) Heatmap of genes for known axonally-trafficked protein genes 1 and 3 days after SNI and SCI in whole DRG or fluorescence-activated cell sorted (FACS) dorsal column (DC) neurons from Thy1YFP16 mice. (SNI (n=3 mice) and SCI (n=4 mice) for C; n=3 mice per group, except n=4 mice for naïve and SCI 1d for D). **G**) Schematic indicating the experimental design for the *in vivo* conditioning experiment. **H**) Representative images of horizontal sections of the dorsal column labeled with SCG10 and dextran-labeled conditioned axons that was injected into the ipsilateral sciatic nerve. **I**) Quantification of **H** indicating the ratio of mean SCG10 staining intensity between conditioned and unconditioned axons of DC neurons (n=3 mice; One-sample t-test and Wilcoxon signed rank test).

Palmitic acid generated by FASN is also critical for protein palmitoylation ^51^. Both Gap43 and SCG10 proteins are palmitoylated and anterogradely transported after nerve injury ^52^. Palmitoylation of SCG10 facilitates its targeting to the growth cone ^53,54^ and axonal transport of SCG10 is widely used to label regenerating axons ^47^. We found that SNI, but not SCI, induces upregulation of Gap43 and SCG10 at the gene expression level (**Fig 4E-F**). We next tested if SGC10 accumulation at injured axon tips fails to occur in injured ascending sensory axons after SCI. We performed a unilateral conditioning SNI, followed by thoracic SCI along with tracer injection into the conditioned sciatic nerve 3 days later, and histology assessment of the ascending sensory 1d later (**Fig 4G, S5**). Since the tracer labels only ipsilateral dorsal column axons, we can assess both conditioned (ipsilateral, tracer-labeled) and unconditioned (contralateral, unlabeled) axons in the same animal and tissue section. We found that SCG10 accumulates in injured spinal axons of ascending sensory neurons only after a conditioning injury (**Fig 4H-I**). This is consistent with previous findings showing that CNS axons increase axonal transport after peripheral conditioning ^55^. SCG10 accumulation in PNS axon tips is associated with neurite growth and regeneration ^20,56,57^, whereas SCG10 degradation has surprisingly been suggested to promote axon growth after SCI ^58^. Our results reveal that axonal transport of SCG10 fails to occur in ascending sensory neurons after SCI. Whether this impaired transport results in part from decreased protein palmitoylation as a consequence of decreased FASN expression remains to be tested. Together, these results supports the notion that decreased expression of lipid metabolism-related genes after SCI, including FASN, restricts axon regenerative capacity after SCI.

## Discussion

Our results reveal that ascending sensory neurons respond to SCI and alter their transcriptome in a way that differs from SNI. The upregulation of stress response genes such as ATF3 and Jun suggest that ascending sensory neurons can sense injury after SCI and mount a stress response, but this response is not sufficient to promote a growth state. Our analysis of ATF3 target genes further suggests that in the context of SCI, ATF3 regulates a set of genes that may inhibit ascending sensory neuron growth, whereas in the context of SNI, ATF3 is necessary to transform injured neurons into a regenerative state ^28^. We identified downregulation of lipid biosynthesis related genes in ascending sensory neurons after SCI as one possible mechanism that inhibits axon growth. Pharmacologic inhibition of the enzyme generating fatty acid, FASN, decreases axon growth and regeneration *in vitro*. Our results are consistent with the notion that phospholipid synthesis is required for axon growth in cultured neurons ^21,23,40^ and *in vivo* in retinal ganglion cells ^26^.

Since sensory neurons represent only ~10% of cells within the DRG ^15^, and only a subpopulation (the ascending sensory neurons) are injured by a T9 SCI ^34^, we opted to FACS-sort ascending sensory neurons from YFP16 mice in our analysis. We did find that the FACSeq analysis detected fewer DE genes after both SNI and SCI compared to whole DRG RNA-seq. Differences in methodology may underlie these results, since dissociation and FACS sorting induces an acute axon injury in DRG neurons and sorting 100 YFP+ neurons in triplicate introduces added variability due to random sampling. Nonetheless, our analysis revealed many DE genes (>500) in ascending sensory neurons. This is significantly more than the DE genes observed (<100) in FACS-sorted nociceptors after compression SCI ^14^, which are not directly injured by SCI ^34^. This would suggest that injury to a projection neuron which receives primary input from DRG neurons leads to a much smaller change in gene expression than direct injury to the ascending neuron itself. We also found that fewer DE genes identified in FACS-sorted ascending sensory neurons overlapped with DE genes identified from whole DRG after SCI (13%) compared with SNI (~43%). These differences likely reflect the fact that ascending sensory neurons are only a subpopulation within the DRG.

One important finding was that SCI leads to increased expression of ATF3, a transcription factor known to regulate the pro-generative program after peripheral nerve injury ^28^. However, after SCI, the genes containing an ATF3 binding motif were largely distinct from those after nerve injury. Our analyses also suggest that ATF3 may be responsible for the downregulation of lipid metabolism related genes. These findings may explain why ATF3 overexpression can promote peripheral nerve regeneration ^59^, but fails to do so in several models of CNS injury ^30,60,61^. Changes in epigenomic signatures ^62^ or expression of transcription factors acting as co-factors ^63^ after SCI and SNI may contribute to the difference in expression of ATF3 regulated genes.

Another relevant finding was that the numbers of DE genes in ascending sensory neurons increased from 1 to 3 days after SNI but decreased from 1 to 3 days after SCI. This suggests that the SCI response is not sustained over time and could explain why few transcriptional changes in whole DRG were previously observed 7 days after SCI ^3^. Why transcriptional responses would decrease in time after SCI is unclear, but one possibility is inhibitory signaling from newly established CSPG’s at the injury site, which can be observed as early as 1dpi ^64^. The effect of the environment on the neuronal transcriptional response is supported by the recent observation that a regenerative transcriptome is not sustained in corticospinal tract motor neurons, unless a graft of neural progenitor cells, which promotes robust regeneration, is provided ^65^. Another possibility is anatomical, since SCI leaves the peripheral sensory branch intact. Therefore, the continuous sensory input from the peripheral branch may restrict the expression of an injury response over time. This is consistent with the notion that electrical activity suppresses axon growth and limits regenerative ability in the injured CNS ^66–68^. In addition to leaving the peripheral axon intact, thoracic SCI also leaves the descending axon branch in the spinal cord intact. Indeed, after entering the spinal cord, the central sensory axon bifurcates, with one axon branch ascending and one axon branch descending in the spinal cord ^69^. It has been shown by *in vivo* two-photon imaging that a surviving intact descending branch suppresses regenerative response of the injured ascending branch, while eliminating both leads to a strong regenerative response ^70^. It is thus possible that the spared descending axonal branch after thoracic SCI impairs the expression of a full regeneration program.

Our transcriptional profile analysis from whole DRG and FACS-sorted ascending sensory neurons both suggested a potential role for downregulation of biosynthetic and lipid metabolic pathways after SCI in repressing axon regeneration. Decreased lipid metabolism after SCI in whole DRG is consistent with our recent observation that FASN expression in satellite glial cells promotes axon regeneration in peripheral nerves in part by activating PPARa in satellite glial cells ^43^. Here we found that *Fasn* is also expressed in neurons, and is more significantly downregulated after SCI than SNI in ascending sensory neurons. Using pure neuronal cultures, we found that FASN inhibition decreased neurite outgrowth and axon regeneration *in vitro.* The axon growth defect was rescued by the PPARγ agonist rosiglitazone, suggesting that in neurons, FASN functions in part to generate ligands for PPARγ which promotes neuronal regeneration after injury ^50^. We also observed that SCG10 accumulation at the tip of injured ascending sensory axons is impaired after SCI. This is consistent with the notion that decline of intrinsic axon regenerative ability is associated with selective exclusion of key molecules from CNS axons, and that manipulation of transport can enhance regeneration ^71^. Whether SCG10 decreased transport after SCI is the result of decreased palmitoylation, a key function of FASN ^51^, remains to be tested. FASN is also thought to facilitate membrane outgrowth ^72^, in part by cooperating with protrudin, which is known to induce neurite formation ^73^. Together, these findings indicate that FASN may impact axon growth at multiple levels. Whether FASN expression can increase axon growth after SCI will require future investigations.

## Materials & Methods

### Experimental Animals & Surgical Procedures

All mice were approved by the Washington University School of Medicine Institutional Animal Care and Use Committee (IACUC) under protocol A3381-01. All experiments were performed in accordance with the relevant guidelines and regulations. All experimental protocols involving mice were approved by Washington University School of Medicine (protocol #20180128). Mice were housed and cared for in the Washington University School of Medicine animal care facility. This facility is accredited by the Association for Assessment & Accreditation of Laboratory Animal Care (AALAC) and conforms to the PHS guidelines for Animal Care. Accreditation - 7/18/97, USDA Accreditation: Registration # 43-R-008.

All surgical procedures were performed under isofluorane anesthesia according to approved guidelines by the Washington University in St. Louis School of Medicine Institutional Animal Care and Use Committee. Adult female mice (C57/Bl6, Envigo and Thy1-YFP16, Jackson Laboratory; 10-20 weeks) were used due to ease of bladder voiding after SCI. Buprenorphine SR-LAB (1mg/kg, subcutaneous, ZooPharm) was administered 1 hour before surgery for analgesia. During surgery mice were anesthetized with 2.5% isoflurane. Surgical sites were shaved and disinfected with povidone-iodine solution (Ricca) and alcohol. After surgery underlying tissue was sutured with absorbable sutures and the skin closed with wound clips.

For sciatic nerve injury (SNI), a small skin incision (~1cm) was made at midthigh, the underlying tissue separated by blunt dissection, and the right sciatic nerve exposed and crushed with fine forceps for 5 seconds. For tracer injection into the sciatic nerve, 2ul of 10% dextran, texas red (Thermo Fisher, Cat # D-3328) was injected using a 5ul Hamilton syringe. For spinal cord injury (SCI), a small midline skin incision (~1cm) was made over the thoracic vertebrae at T9-T10, the paraspinal muscles freed, and the vertebral column stabilized with metal clamps placed under the T9/10 transverse processes. Dorsal laminectomy at T9/10 was performed with laminectomy forceps, the dura removed with fine forceps, and the dorsal column transversely cut using fine iridectomy scissors.

### Tissue Processing & Immunohistochemistry

Immediately after euthanasia mice were transcardially perfused with 10mL of PBS followed by 10mL of 4% paraformaldehyde (PFA) in PBS (FD Neurotechnologies; PF101). Tissue was dissected and post-fixed in 4% PFA for 4 hours and then cryoprotected in 30% sucrose overnight. All tissue was sectioned using a cryostat at 10um except for spinal tissue (20um). For immunohistochemistry, tissue sections were blocked in 5% donkey serum (Sigma-Aldrich; Catalog #D9663) in 0.2% PBST (1hr) and incubated overnight at 4°C in primary antibodies diluted in the blocking solution. The next day sections were incubated in secondary antibodies (1:500; 1hr) in PBS and coverslipped in ProLong Gold antifade mounting media (Thermo Fisher). Between steps tissue sections were washed with PBS. For image acquisition, a Nikon TE-2000E microscope equipped with a Prior ProScan3 motorized stage was used. Nikon Elements software was used for image analysis.

### Cell Culture

For adult DRG cultures, mice were transcardially perfused immediately after euthanasia with 10mL of Hanks’ balanced salt solution with 10 mM HEPES (HBSS-H) and L4 DRG dissected into HBSS-H on ice. DRG were treated with papain (15 U/ml, Worthington Biochemical) and collagenase (1.5 mg/ml, Sigma-Aldrich) in HBSS-H at 37C for 20 minutes, then dissociated by trituration into 3 or 4 wells of a 24-well glass-bottom plate coated with 100 μg/ml poly-D-lysine and 3 μg/ml laminin. Culture media consisted of Neurobasal-A medium, B27 plus, glutaMAX, and penicillin/streptomycin. For experiments assessing Fasn inhibition, vehicle (0.05% DMSO) or platensimycin (1μm) was added to the media 30 minutes after plating. After 24hr cells were fixed with 4% PFA for 20 minutes and then immunostained as described above.

For *in vitro* regeneration assay with embryonic DRG cultures, DRG were isolated from time pregnant e13.5 CD-1 mice. DRG were kept in cold HBSS media until all DRGs were collected. After a short centrifugation, dissection media was aspirated and ganglia were digested in .05% Trypsin-EDTA for 25 minutes in 37°C. Next, cells were pelleted by centrifuging for 2 minutes at 500 x g, the supernatant was aspirated, and Neurobasal was added. Cells were then triturated 25x and added to the growth medium containing Neurobasal media, B27 Plus, 1ng/ml NGF, Glutamax and Pen/Strep, with 5μM 5-deoxyfluoruridine (FDU). Approximately 10,000 cells were added to each well in a 2.5 μl spot. Spotted cells were allowed to adhere for 10 minutes before the addition of the growth medium. Plates were pre-coated with 100μg/ml poly-D-lysine. Vehicle (0.05% DMSO), platensimycin (1μM) (gift from Merck Research Labs) or rosiglitazone (10μM) (Sigma-Aldrich; Catalog # R2408) was added to culture on DIV6. Cells were then injured using an 8mm microtome blade on DIV7 and fixed 24h later. Cells were washed with PBS and stained for SCG10 as described above.

### Image Analysis

Total neurons were identified using TUJ1 (BioLegend, Cat # 801202, 1:1000) or for neuronal nuclei with Islet-1 (Novus Biologicals, Cat # af1837-sp, 1:500) antibodies, while YFP positive neurons were identified by endogenous GFP expression. Nuclear expression of ATF3 (Novus Biologicals, Cat # NBP1-85816, 1:250) and Jun (Cell Signaling, Cat # 9165S, 1:500) was assessed in all Islet-1 positive neuronal nuclei. NF200 positive (Abcam, Cat # ab4680, 1:1000) and TrkA negative (EMD Millipore, Cat # 06-574, 1:500) staining was used to characterize ascending sensory neurons in YFP positive neurons. For cell culture experiments in Thy1 YFP16 mice, percent axon initiation (neurons with axon length > 50um) and radial length (distance from soma to furthest axonal point) was manually measured for TUJ1 positive and YFP positive neurons. For FASN inhibition experiments in adult DRG cultures, automated cell body counts and neurite tracing and length quantification was performed using Nikon elements analysis explorer software. For embryonic dorsal root ganglia experiments, regenerative length was measured from the visible blade mark to the end of the regenerating axons. Each technical replicate was measured 4-6 times and three technical replicates were measured per biological replicate. To assess ipsilateral versus contralateral SCG10 levels in the dorsal column, a rectangle was first drawn around the ipsilateral dextran-labeled conditioned axons (up to 1mm away from the injury site), the image channel switched to show SCG10 labeling, and mean pixel intensity (SCG10 staining) measured using Nikon elements software. The rectangle was then moved to the immediate contralateral side of the dorsal column and SCG10 labeling similarly measured.

### Whole DRG RNA sequencing

L4 DRG were dissected and homogenized in 300μl of lysis buffer on ice and RNA purified using the PureLink RNA Mini kit (Thermo Fisher), which was submitted to the Genome Technology Access Center at Washington University for library preparation and sequencing. RNA quality was assessed using an Agilent Bioanalyzer (260/280 > 1.9; RIN > 8.0). Samples were subjected to DNase treatment. rRNA depletion was achieved with the Ribo-Zero rRNA removal kit. Library preparation was performed using the SMARTer kit (Clontech), and sequencing performed on an Illumina HiSeq3000.

Briefly, sequences are adapter-trimmed using Cutadapt ^74^ and subjected to quality control using PRINSEQ ^75^ and aligned to mouse genome GRCm38/mm10 using STAR ^76^. Sequencing performance was assessed for total number of aligned reads, total number of uniquely aligned reads, genes and transcripts detected, ribosomal fraction, known junction saturation, and reads distribution over known gene models with RSeQC ^77^. Reads in features were counted using HTseq ^78^. Genes differentially expressed between conditions were identified using DESeq2 with a false discovery rate (FDR) adjusted p values < 0.1, which includes a Benjamini-Hochberg correction ^79^. Variance stabilizing transformation (VST) normalized counts were calculated using DESeq2, and normalized gene counts were converted to Z scores for plotting. Heatmaps were generated using heatmap.2 function of the gplots R package.

### Fluorescence-Activated Cell Sorting RNA Sequencing (FACS-seq)

L4 DRG from Thy1-YFP16 mice were dissociated as described above and FACS-sorted as previously described ^36^. Briefly, after dissociation cells were passed through a 70μm cell strainer and resuspended in PBS with 2% fetal calf serum. L4 DRG cells were sorted by GFP signal in triplicate for each sample (100 cells per well; each sample of 100 cells was obtained from one mouse) and submitted to the Genome Technology Access Center at Washington University for library preparation and sequencing. Library preparation was performed using the SMARTer Ultra LowRNA kit (Clontech) and sequencing performed on an Illumina HiSeq3000. RNA quality was assessed using an Agilent Bioanalyzer (260/280 > 1.9; RIN > 8.0).

Differentially expressed gene analysis were performed as described above except noted below. Genes with < 20 reads in all samples were excluded from further analysis. Outlier samples were identified using robust principal component analysis according to the ROBPCA, and GRID algorithms implemented in rrcov R package and were removed from further analysis ^80^. The factors of unwanted variation were estimated using RUVr with k = 2 and were modeled in the DESeq2 design formula ^81^.

### ATF3 motif analysis

ATF3 position frequency matrix (MA0605.1) were obtained from JASPAR database (http://jaspar.genereg.net/). The Patser program calculates the probability of observing a sequence with a particular score or greater ^38,82^ for the given matrix and determines the default cutoff score based on that P-value. Sequence 5kb upstream of ATG start codon of a given gene was scanned using Patser to identify ATF3 binding sites. Any gene with at least one ATF3 binding site was counted as ATF3 target gene. R package clusterProfiler ^83^ was used for GO and KEGG pathway enrichment analysis and plotting. GO and KEGG pathway terms with FDR corrected P value < 0.05 were considered as significant.

### Data Deposition

FASTQ files were deposited at the NCBI GEO database. Accession: GSE149646

Reviewer token: otqlcagytrmdryd

### Statistical Analysis

All quantifications used for statistical analysis was performed by experimenters blinded to treatment conditions. DESeq2 with a false discovery rate (FDR) adjusted p values < 0.1. GraphPad Prism software was used for statistical analysis. Statistical tests, sample sizes, and p-values are reported in the legend for each figure. Statistical significance was defined as p < 0.05 or adjusted p < 0.1 for RNA-seq experiments. Error bars indicate the standard error of the mean (SEM).

## Acknowledgments

We would like to thank members of the Cavalli lab for valuable discussions. We thank Clay Semenkovich for helpful discussions on fatty acid synthesis and for providing us with the FASN inhibitor platensimycin. We also thank Anushree Seth and Madison Mack in association with InPrint for illustration in Fig.1a and 2a. This work was funded in part by a post-doctoral fellowship from the Craig H. Neilsen Foundation to E.E.E, by NIH grant NS096034, NS082446 and NS111719 to V.C.

## Author contribution

EEE and VC designed experiments. EEE performed and analyzed experiments. OA performed *in vitro* DRG regeneration experiments. DC performed FACS of YFP positive neurons. TMG and GZ analyzed RNAseq data. EEE and VC wrote the manuscript. All authors reviewed the manuscript.

## Supplementary information

### Supplementary Figure legends

**Figure S1.**
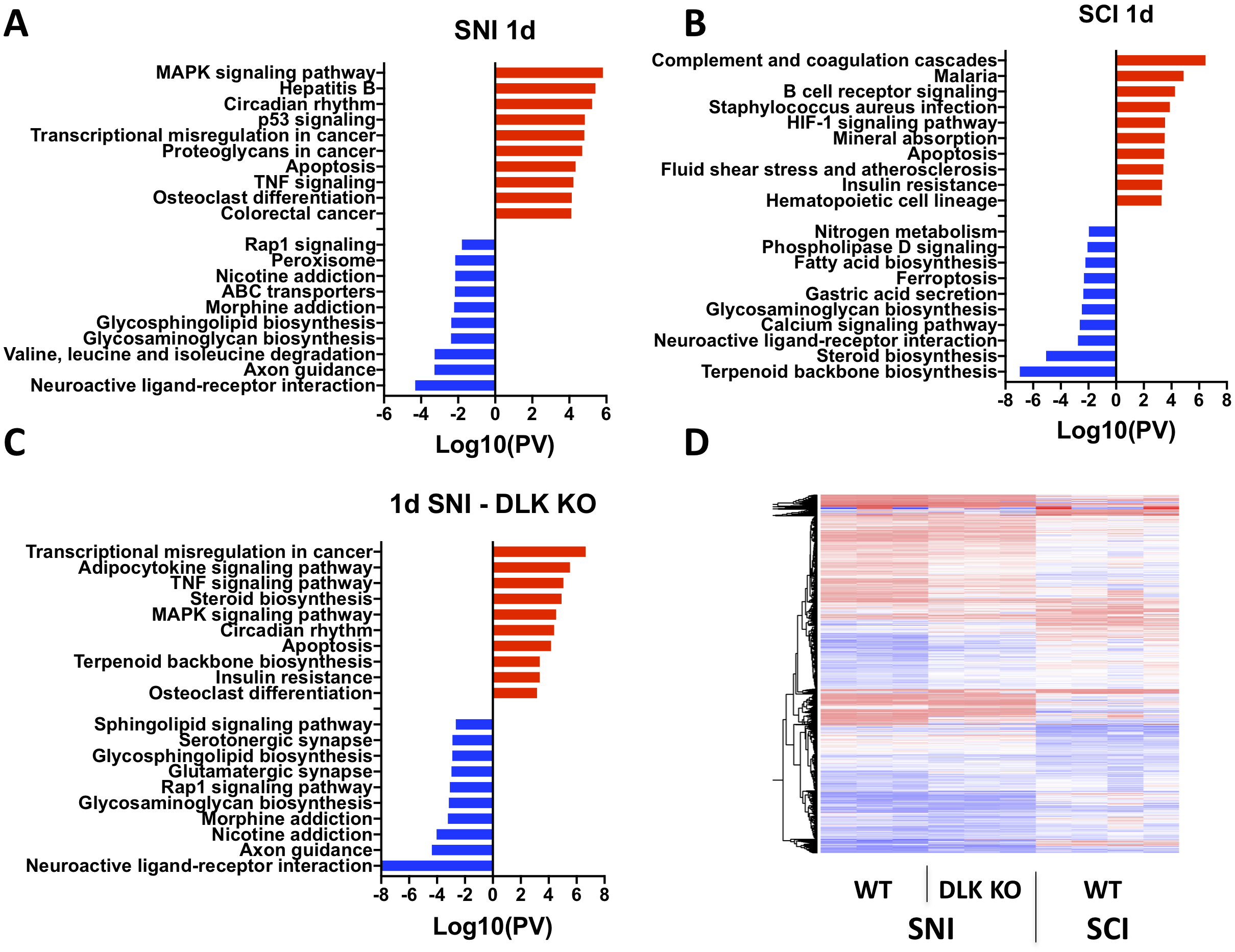
Pathway analysis and transcriptional comparison after sciatic nerve injury (SNI) and spinal cord injury (SCI) in dorsal root ganglion (DRG). **A-C**) Pathway analysis for the most significantly enriched pathways associated with upregulated (red) and downregulated (blue) differentially expressed (DE) genes after SNI (n=3 mice) or SCI (n=4 mice) in wildtype or SNI in dual leucine zipper kinase (DLK) knockout mice (n=3 mice; p-adj < 0.1; KEGG 2016). **D**) Heatmap of all upregulated (red) and downregulated (blue) DE genes for each subject and each condition.

**Figure S2.**
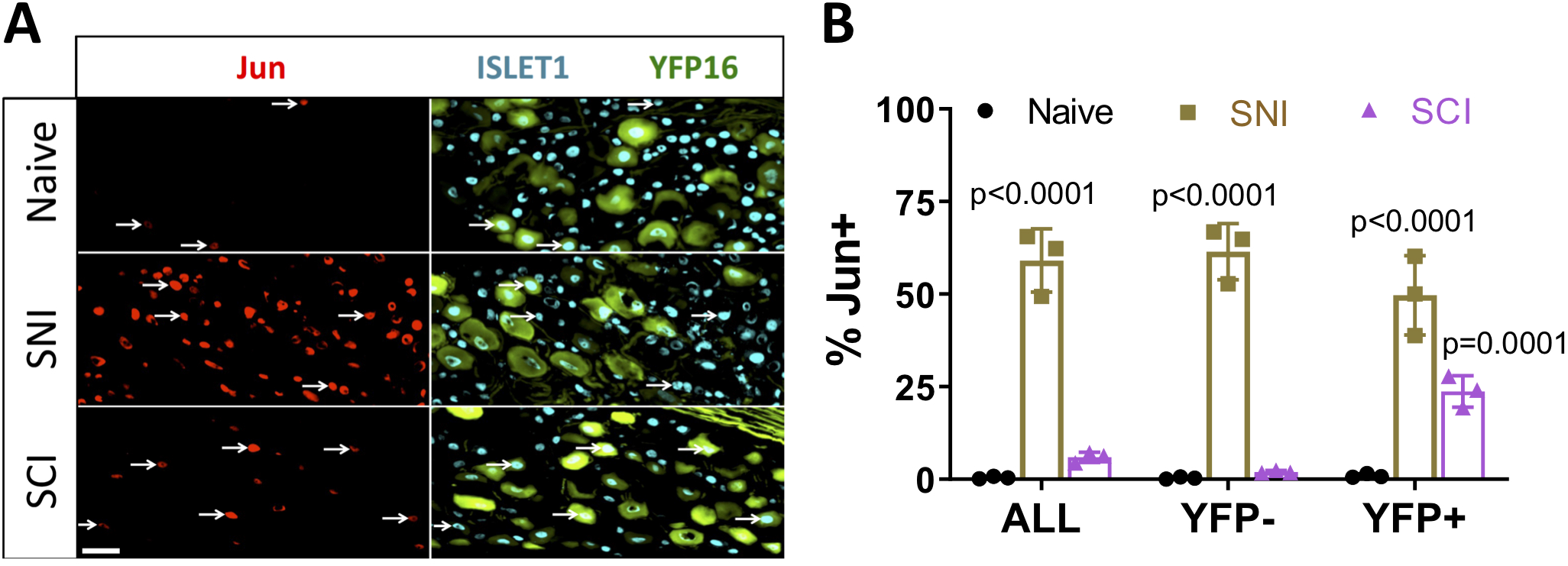
Jun upregulation is specific to dorsal column (DC) neurons after spinal cord injury (SCI). **A**) Representative images of L4 dorsal root ganglion (DRG) neurons labeled with Jun and Islet1 antibodies in Thy1YFP16 mice in naive or 3 days after sciatic nerve injury (SNI) or SCI. **B**) Quantification of **A** indicating percentage of Jun positive, Islet-1 labeled neuronal nuclei in all neurons, as well as YFP negative and YFP positive neurons, for each condition (n=3 mice/group; 2-way ANOVA). White arrowheads point to Jun positive neuronal nuclei. *p < 0.05, ***p < 0.001

**Figure S3.**
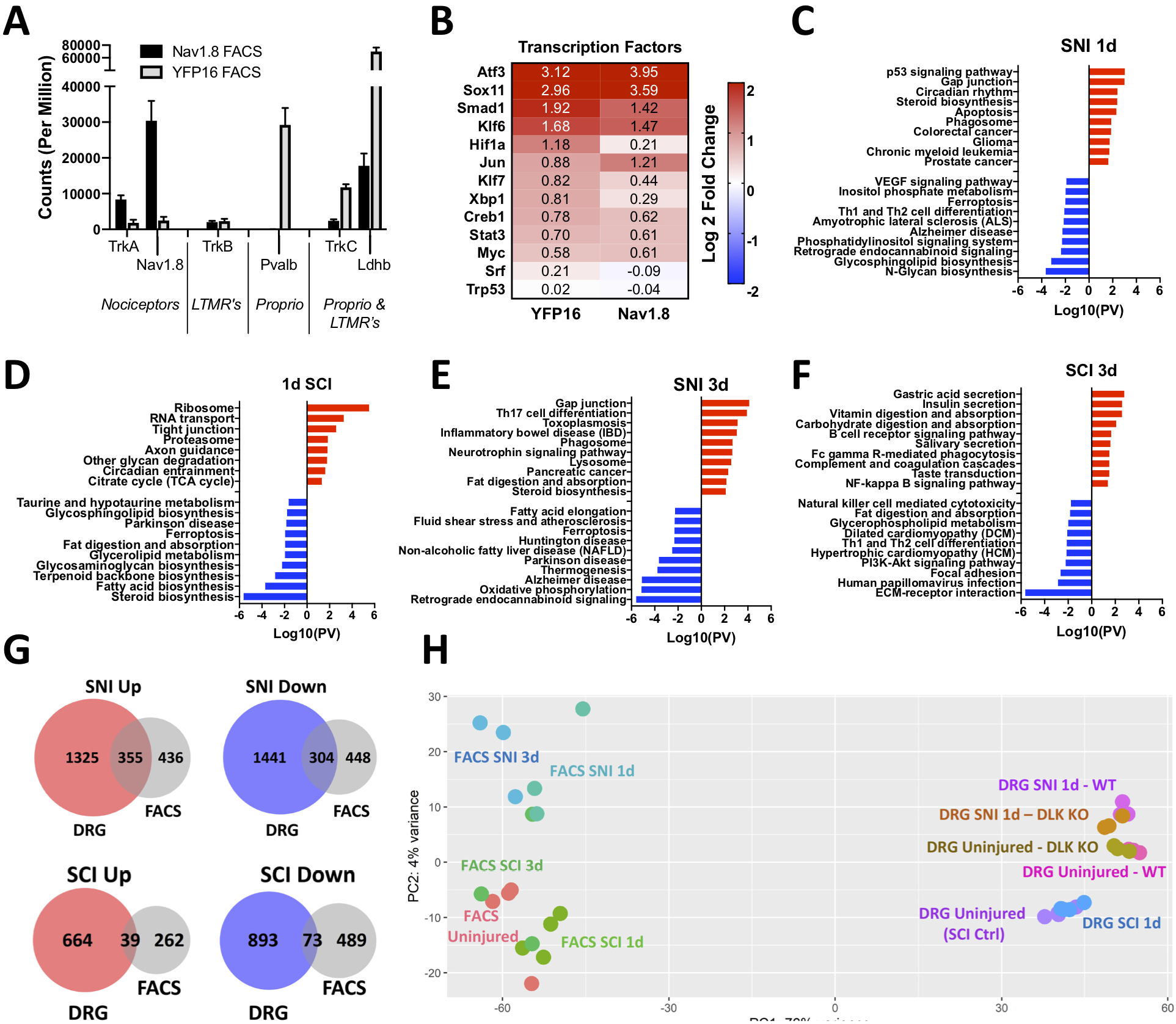
Pathway analysis and transcriptional response of DC neurons 1 and 3 days after sciatic nerve injury (SNI) and spinal cord injury (SCI). **A**) Comparison of total counts of genes associated with three predominant neuronal subtypes in the DRG (nociceptors, low-threshold mechanoreceptors (LTMR’s), and proprioceptors (Proprio)) between fluorescence-activated cell sorted (FACS) nociceptors (Nav1.8) and DC neurons (Thy1YFP16). **B**) Heatmap of known regeneration-associated transcription factors (RATF’s) 3 days after SNI in FACS sorted nociceptors (Nav1.8) and DC neurons (Thy1YFP16). **C-F**) Pathway analysis for the most significantly enriched pathways associated with upregulated (red) and downregulated (blue) differentially expressed (DE) genes 1 and 3 days after SNI (n=3 mice) or SCI (n=4 mice (1d), n=3 mice (3d)) in DC neurons (KEGG 2016). **G-H**) Proportional venn diagrams for DE genes upregulated (red) or downregulated (blue) 1 day after SNI and SCI in whole DRG (p-adj<0.1) compared with DC neurons (p-adj<0.1, RUvR = 2).

**Figure S4.**
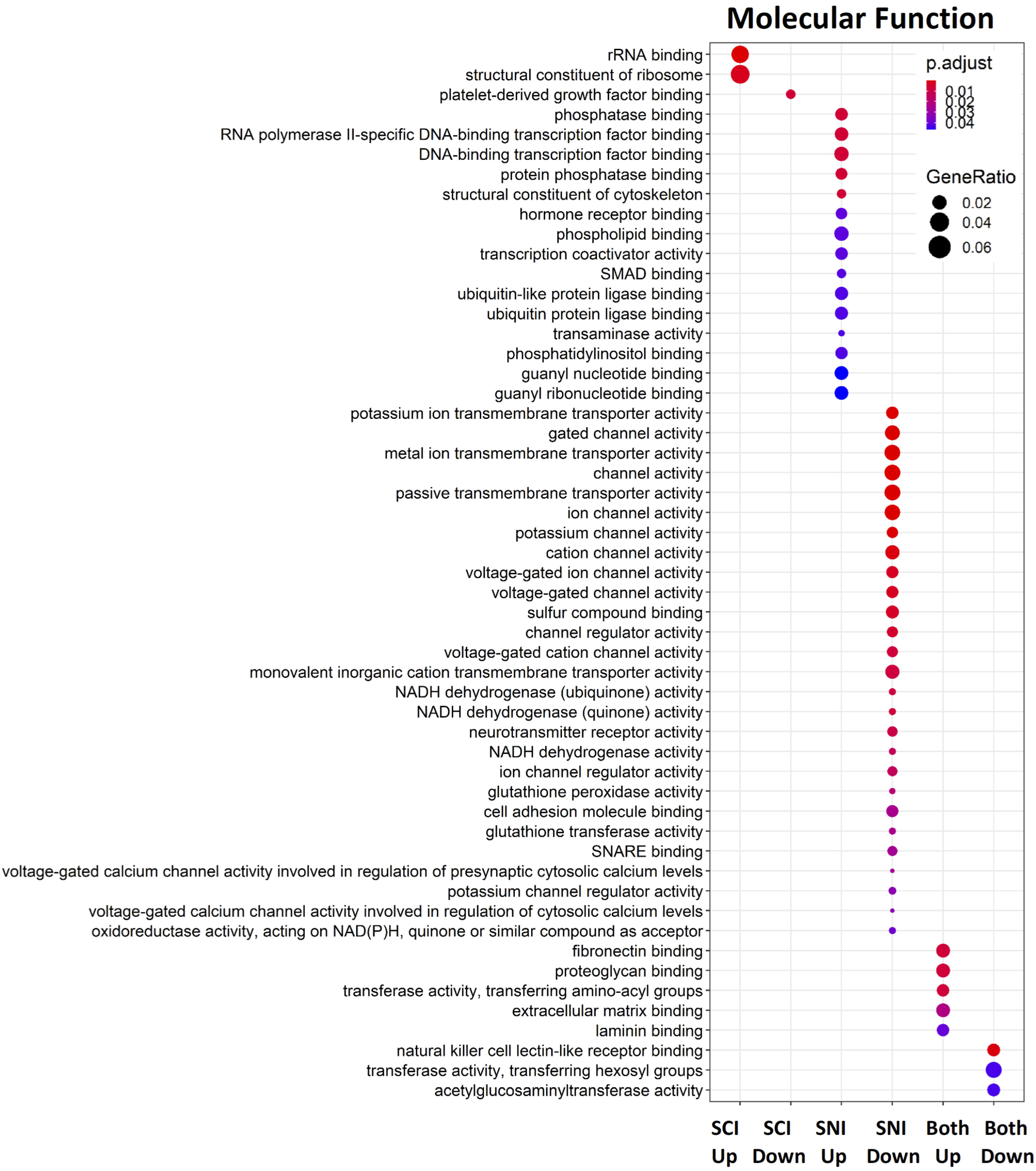
GO analysis of genes containing and ATF3 binding motif after SNI or SCI. Gene Ontology (GO) enrichment analysis was performed on differentially expressed (DE) genes that have ATF3 binding sites in the promoter upstream sequences. DE genes from 1d and 3d post injured were combined in each injury condition. The DE genes were categorized into genes that were uniquely upregulated in SCI or SNI, uniquely downregulated in SCI or SNI as well as genes that were up- or down-regulated in both SCI and SNI. The graph shows the GO terms in the Molecular Function category enriched in each gene list.

**Figure S5.**
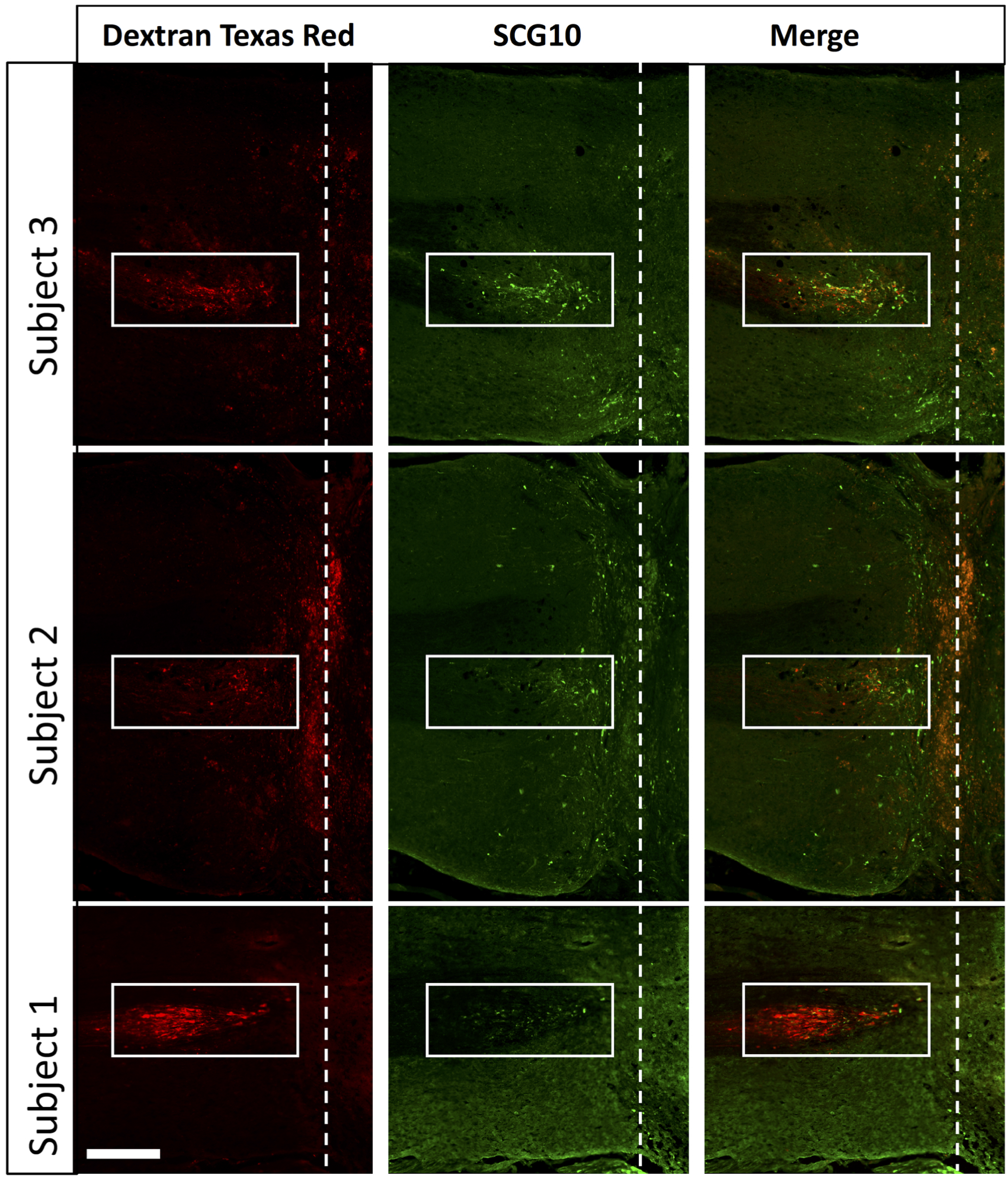
Conditioning nerve injury leads to accumulation of SCG10 in injured ascending sensory axons. Horizontal sections of the dorsal column labeled with SCG10 and dextran-labeled conditioned axons that was injected into the ipsilateral sciatic nerve. Representative horizontal spinal images are shown for 3 different subjects.

### Supplementary Tables

**Table 1.** List of genes significantly upregulated after SCI (green) or SNI in wildtype (gray) and DLK KO (blue) mice 1d after injury in whole DRG.

**Table 2**. List of genes significantly downregulated after SCI (green) or SNI in wildtype (gray) and DLK KO (blue) mice 1d after injury in whole DRG.

**Table 3**. List of all genes after SCI (green) or SNI in wildtype (gray) and DLK KO (blue) mice 1d after injury in whole DRG.

**Table 4**. List of genes that were inversely expressed between SNI and SCI 1d after injury in whole DRG.

**Table 5**. List of all genes after SCI (green) or SNI in mice 1d (lighter shade) or 3d (darker shade) after injury in FACS-sorted dorsal column neurons.

**Table 6**. List of genes significantly upregulated after SCI (green) or SNI in mice 1d (lighter shade) or 3d (darker shade) after injury in FACS-sorted dorsal column neurons.

**Table 7**. List of genes significantly downregulated after SCI (green) or SNI in mice 1d (lighter shade) or 3d (darker shade) after injury in FACS-sorted dorsal column neurons.

**Table 8.** List of genes containing ATF3 binding motifs regulated by SNI and SCI

